# Alternative splicing switches STIM1 targeting to specialized membrane contact sites and modifies SOCE

**DOI:** 10.1101/2020.03.25.005199

**Authors:** Mona L. Knapp, Kathrin Förderer, Dalia Alansary, Martin Jung, Yvonne Schwarz, Annette Lis, Barbara A. Niemeyer

## Abstract

Alternative splicing is a potent modifier of protein function. Stromal interaction molecule 1 (Stim1) is the essential activator molecule of store-operated Ca^2+^ entry (SOCE) and a sorting regulator of certain ER proteins such as Stimulator of interferon genes (STING). Here, we characterize a conserved new variant, Stim1A, where splice-insertion translates into an additional C-terminal domain. We find prominent expression of *exonA* mRNA in testes, astrocytes, kidney and heart and confirm Stim1A protein in Western blot of testes. In situ, endogenous Stim1 with domain A, but not Stim1 without domain A localizes to unique adhesion junctions and to specialized membrane retrieval sites (tubulobulbar complexes) in testes. Functionally, using Ca^2+^ imaging and patch-clamp analysis, Stim1A shows a dominant-negative effect on SOCE and I_CRAC_, despite normal clustering and interaction with Orai1 investigated by combined TIRF and FRET analyses and as confirmed by an increased SOCE upon knock-down of endogenous Stim1A in astrocytes. Mutational analyses in conjunction with imaging and patch-clamp analyses of residues either in domain A or within the N-terminal region of Orai1 demonstrate a specific defect in stabilized channel gating. Our findings demonstrate that cell-type specific splicing of STIM1 adds both an intracellular targeting switch and adapts SOCE to meet the Ca^2+^ requirements of specific subcellular contact sites.

## Introduction

STIM and ORAI proteins are the hallmark constituents of store-operated Ca^2+^ entry (SOCE), critically important for T cell function, but have since been found in virtually all cell types. Especially relatively slow and long-lasting processes such as gene expression and cell proliferation cells rely on SOCE ^1,2^. SOCE is triggered by receptor mediated release of Ca^2+^ from the endoplasmic reticulum (ER), which is sensed by STIM1 and STIM2 proteins through their respective luminal EF hands which then cluster and trap ORAI channels located in the plasma membrane (PM) into ER-PM junctions (reviewed in ^3–6^). In patients and murine gene-modified models, loss-of-function mutations (LOF) of either STIM1 or ORAI1 lead to severe immune pathologies, myopathies, defects in enamel formation and anhidrosis (reviewed in ^7–9^). In mice, global deletion of *Stim1* results in a severe reduction in the number of offspring and those that are born die within the first weeks ^10^. The early lethality indicates essential roles for Stim1 that are independent of their immune cell function as also indicated by the fact that also gain-of-function (GOF) mutations result in multisystemic phenotypes (see above). A mouse model expressing the STIM1 gain-of-function mutation R304W, which causes Stormorken syndrome in humans, also shows very few surviving offspring with small size, abnormal bone architecture, abnormal epithelial cell fate as well as defects in skeletal muscle, spleen and eye ^11,12^. Cardiomyocyte-restricted deletion of STIM1 (crSTIM1-KO) results in age-dependent endoplasmic reticulum stress, altered mitochondrial morphology, and dilated cardiomyopathy ^13^, promotes development of arrythmogenic discordant alternans ^14^ and has also been reported to induce dysregulation of both cardiac glucose and lipid metabolism ^13^. Metabolic reprogramming in the absence of STIM1 and STIM2 has also been shown in T cells whereby STIM proteins control clonal expansion ^15^.

The question we set out to explore was if all of these highly diverse functions of STIM1 are indeed mediated by exactly the same protein, or if and how alternative splicing modifies STIM1 structure and function to adapt to cell-type specific requirements. In the past, we and others have identified a splice variant of STIM2 (STIM2.1; STIM2ß), which prevents interaction and gating of Orai channels ^16,17^ thus acts as a strong dominant-negative regulator of SOCE by altering the density of activated Orai complexes ^18^ and has recently been shown to regulate myogenesis by controlling SOCE dependent transcriptional factors ^19^. Given the existence of this inhibitory splice variant, the often small effects seen upon genetic deletion of *STIM2* may indeed underestimate a STIM2 mediated effect, if the concomitant deletion of expressed *STIM2*.*1* deletes the negative regulator STIM2.1/STIM2ß. The same line of thought might apply to potential splice variants of *STIM1*. Darbellay et al. ^20^ were the first to show that alternative splicing of *STIM1* leads to a longer STIM1 protein, STIM1L, which is expressed exclusively in adult human muscle fibers and in *in-vitro* differentiated myotubes ^20^. Here, splice insertion after exon 11 translates into 106 additional C-terminal residues which contain an actin-binding domain. STIM1L forms permanent clusters with Orai1, allowing faster activation which is especially needed during repetitive stimulation. While Horinouchi and coworkers ^21^ confirmed the existence of STIM1L in skeletal muscle and found it to inhibit TRPC3 and TRPC6 activity, Luo et al. ^22^ found STIM1L protein also in neonatal rat cardiomyocytes, with decreasing expression during postnatal cardiac development. Upon agonist- or afterload-induced cardiac stress, STIM1L expression re-emerged. Sauc et al. ^23^ investigated the molecular mechanism of STIM1 vs. STIM1L gating in more detail. While STIM1 was able to expand cortical ER cisternae, STIM1L was unable to do so, but could trap and gate Orai1 channels without remodeling cortical ER ^23^. These results already highlight the importance of understanding *STIM* splicing.

In the current work, we identify several additional STIM1 splice variants with distinct mRNA expression and differential subcellular targeting to highly specialized ER contact sites as shown on protein level by differentiating antibodies. We focus on the functional characterization of a full length variant containing an alternative exon, namely *exon 11 (A)*, which translates into an in frame 31 amino acid domain A. Stim1A is a dominant negative regulator of SOCE in astrocytes and upon heterologous expression and biophysical analysis by imaging, FRET and patch-clamp in conjunction with mutagenesis shows that insertion of domain A prevents full gating of ORAI channels in a sequence specific manner. Furthermore, its subcellular targeting points towards a role of Domain A containing Stim1 in membrane retrieval.

## Results

### Stim1 *exon 11 (A)* is an evolutionary conserved expressed Stim1 splice variant

Both *Stim1* and *Stim2* are genes containing multiple exons and database mining predicts several splice variants with three predicted murine variants (NM_001374058.1, XM_006507535.4, XM_006507536.2) and human EST clones (BQ068737.1, BU553936.1) containing an additional *exon 11 (A)* insertion between *exon 10* and *exon 12* (formerly termed exon 11). The *exon 12* and *exon 14* boundaries harbor mutually exclusive alternative *exons 13 (L)* or *13 (B)* (Fig. 1a). *Exon A* splicing extends the C-terminal domain by 93 nucleotides and translates to a 31 amino acid in-frame insertion (domain A) downstream of C-terminal residue 491. Using cDNA derived from different murine tissues, we initially devised both a flanking PCR as well as qRT-PCR strategies to screen for the presence of the alternative *exon A* versus expression of the conventional variant (*ØA=wt*) (Fig. 1a). This strategy enabled us to detect multiple splice events because primers extend over several splice sites and could result in 310 bp for *Stim1 wt*, 402 bp for *exon A*, 438 bp for *exon AB*, 613 bp for *exon L* and 346 bp for an *exon B* splice event. We were able to detect 310 bp (*wt*) products in all selected tissues as well as a prominent additional 402 bp (*exon A*) product in testes and astrocytes. Only in skeletal muscle we detected the 613 bp product (*exon L*) (Fig. 1c). Using quantitative real time PCR either with an exon A specific primer or a primer bridging the *exon 10/exon 11* boundary (*ØA*) (Fig. 1b), we confirmed significant relative expression of *exon A* in testes, heart, astrocytes and kidney, while CD8^+^ T cells did not show detectable expression of *exon A* (Fig. 1d). The 2^−(ΔCq)^ values shown in Supplementary Figure 1b demonstrate that expression of *exon A* is seen in most tissues with less variability compared to *Stim1ØA* expression. As database mining also indicated that expression of *exon A* may occur in conjunction with alternative splicing within the *5’UTR* region (putative transcript XM_006507535.4), we confirmed the existence of *exon A* (reverse primer) with the regular STIM1 N-term by PCR using forward primers specific for the *5’UTR (wt), 5’UTR (alt 1)* or *5’UTR (alt 2)* (Fig. 1a and S1a). Both *5’UTR* splice events would lead to deletion of *exon 1* and result in translation at the first *ATG* within *exon 2*. Translation of these variants would lead to proteins lacking the signal peptide and essential parts of the first EF hand to start at M75 of Stim1 and are not functionally investigated here. Supplementary Figure 1c shows a prominent PCR product of the *5’UTR (wt)* together with an *A*-specific reverse primer, a weaker product of *A* with the *alt1* primer (26% of *A* products) and no product of *A* with the *alt2* primer in testes. Using the *non-A* reverse primer, we also detect *5’UTR (wt)* as well as *alt1* (5% of *ØA* products) PCR products (Fig. S1c). We also tested a reverse primer annealing within alternate *exon 13 (B)*, which resulted in a very minor product, potentially resembling an *exon AB* splice combination. To quantitate the relative amount of N-terminal splicing, we also used qRT-PCR primers together with a reverse primer within *exon 2* (Fig. S1a) to show that in testes, 86% of all *Stim1* variants contain the wt 5’UTR (Fig. S1d). Having confirmed the existence of *exon A* within *Stim1*, we cloned the entire open reading frame of murine *Stim1A* from cDNA into bicistronic- or directly tagged expression vectors and also inserted *exon A* into human *STIM1*. The resulting constructs translate to a 716 amino acid protein in which domain A is in close proximity to the negatively charged inhibitory domain (ID) (Fig. 1e). 71% of the amino acids encoded by *exon A* are evolutionarily highly conserved from fish, amphibians, reptiles and birds to mammals (Fig. 1f). Only a four amino acid insertion in the middle of domain A appears to have co-evolved with the subclass *Theria* as it is absent in *Monotremes* (egg-laying mammalia, such as *Ornithorhynchus anatinus* = platypus), but present in live bearing mammals (*Theriiformes*, such as *Phascoloarctos cinereus* = koala). Insertion of domain A adds 3,55 kDa to the molecular mass of Stim1, a difference that can only be detected by running optimally separating SDS-PAGE as shown after heterologous expression of both variants (Fig. 1g). Note that insertion of domain A does not significantly alter protein levels as quantified in the right panel of Fig. 1g. Protein extracts from testes show native protein species which run at similar masses as Stim1A and as Stim1 (Fig. 1h). We were also able to detect a similar double band in heart extracts, although longer exposures were necessary (Fig. S1e).

**Figure 1.**
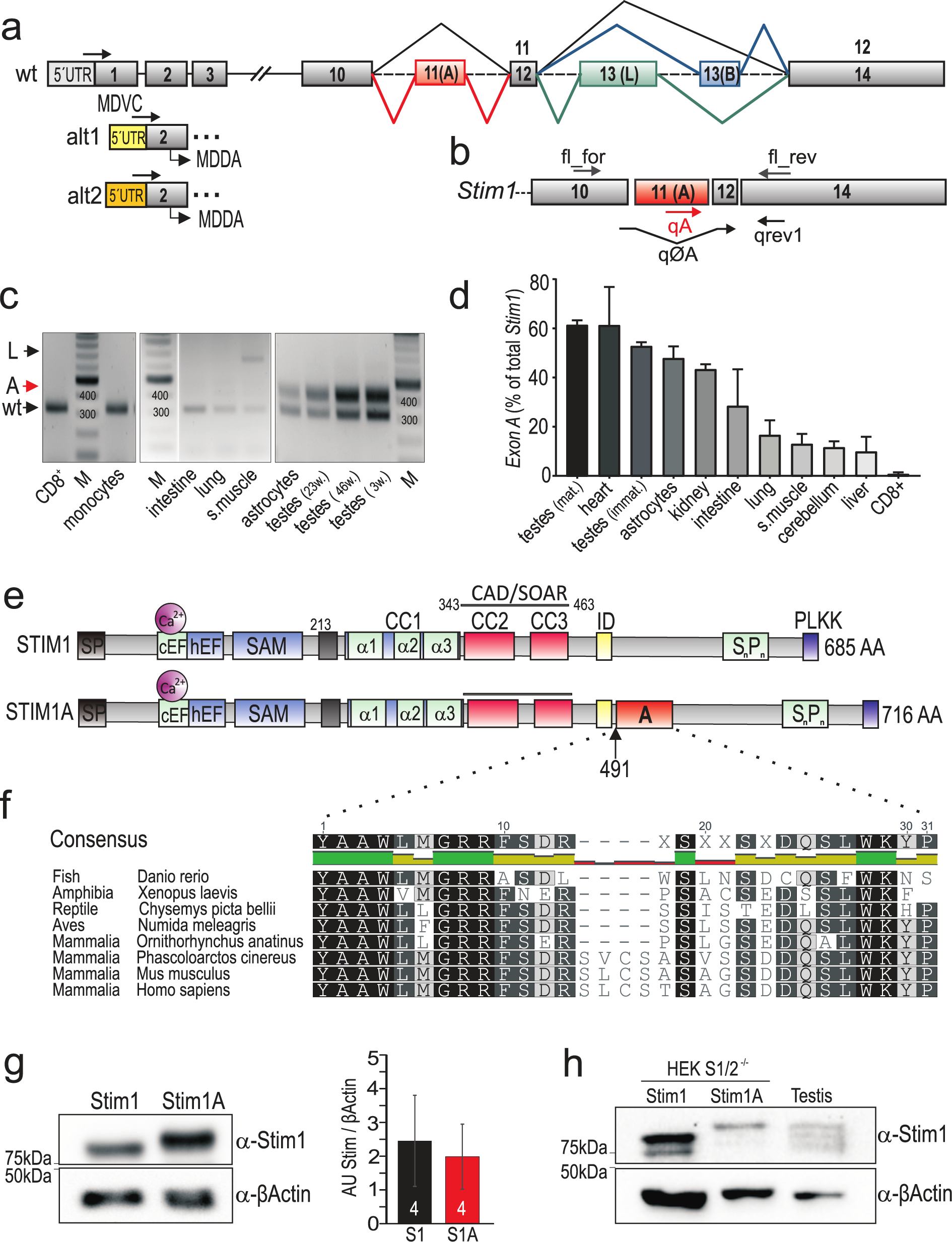
Identification of a novel STIM1 splice variant. (a) Schematic exon structure of *Stim1 gene* with conventional exons depicted in grey. Highlighted in red is the alternative exon 11 (A), and additional mutually exclusive splicing of exon 13 (L, green) or exon 13 (B, blue). Alternative usage of 5’UTR regions are indicated with alt1 and alt2. (b) Annealing sites of primers on the coding mRNA (c) Analytic PCR with primers flanking the splice site using cDNA’s of tissues or cells indicated. (d) Relative expression of *exon A* to total *Stim1* (± SD) in tissues using cDNA derived from 3-4 mice. (e) Schematic protein structure of STIM1 displaying functional motives. Domain A (red) is located 8 amino acids downstream of the inhibitory domain (ID, yellow) after aa 491. (f) Evolutional conservation of domain A. Identical amino acids within black boxes. Western blot showing Stim1 and Stim1A following heterologous expression in HEK293, right panel: quantification from 4 transfections. (h) Heterologous and endogenous levels of Stim1 in testes, using an antibody detecting the N-term of STIM1.

### Domain A targets STIM1 to specialized ER contact sites

In HEK cells, STIM1A does not show an overt localization difference compared to STIM1, which may be due to the lack of specialized- or cell-type specific ER domains. In testes, Sertoli cells “nurse” the developing spermatids and show an intricate and complex ER with several different highly specialized ER domains ^24^. As Sertoli cells are a major cell-type containing Stim1^25,26^ we set out to investigate localization of domain A in murine testes and developed a domain A specific antibody. Specificity compared to Stim1 was confirmed by Western Blots and in IHC using knock-out cell lines expressing either variant (Fig. S2); in addition, we found that the commercially available antibody (H-180) directed against aa 441-620 of STIM1^27^ did not recognize Stim1A and will hence be termed ØA-antibody (Fig. S2). In serial sections of adult mouse testes, we reproduced the STIM1 distribution seen by Vogl et al. in Sertoli cells ^27^ using a polyclonal antibody against the N-terminal domain of STIM1 (Fig. 2b, for no-primary antibody controls see Fig. S3). We then applied the domain A specific antibody in conjunction with the monoclonal ØA antibody (Fig. 2a,c). The overview (Fig. 2a) shows the cross section of a seminiferous tubule. Already at low resolution (Fig. 2a), we noticed intense punctate immunoreactivity with the A antibody in apical regions of Sertoli cells, where spermatids are released and the ER forms specialized contacts to the plasma membrane within the so called tubulobulbar complexes (TBC). TBCs are unique to Sertoli cells and are clathrin-dependent subcellular structures required for internalization of intra-cellular junctions and membranes once spermatids are released ^24,27,28^. At higher magnification, we observed that these punctate structures often contain a less stained lumen (arrows in Fig. 2c, for controls see Fig. S2c). These structures are not recognized by the ØA-antibody, which showed a partially clustered staining within the general ER (Fig. 2c, arrows, ØS1A staining). In basal regions of Sertoli cells, the anti-A antibody labelled contact sites that surround germ cells at different stages (Fig. 2d, see also arrows). These same regions are also recognized by a monoclonal Stim1 antibody directed against the very C-terminal region (Fig. 2d, red panel). To confirm that the globular structures detected by anti-A represent apical TBCs with ER adjacent to tubules of plasma membrane, we co-stained with a monoclonal anti-Orai1 antibody, as Orai1 is the major Orai homolog expressed in testes (Fig. S1f). The overview of Orai1 and domain A staining is shown in Figure 2e. Prominent Orai1 localization can also be observed in punctae within the apical regions of Sertoli cells as well as in interstitial Leydig cells. High resolution imaging within these processes reveal intense labelling of tubular structures (arrows in Fig. 2f, red channel) with adjacent ER structures positive for domain A (arrows in Fig. 2f, green channel). Spermatid heads in this region show highly condensed chromatin, indicative of imminent release to the lumen of the ductus. In contrast to the intense staining seen within the TBC’s, little Orai1 immuno-reactivity is detected at ES regions (Fig. 2g). Summarized in Fig. 2h we depict the observed distribution of Stim1 variants containing domain A in a schematic Sertoli cell (see discussion).

**Figure 2.**
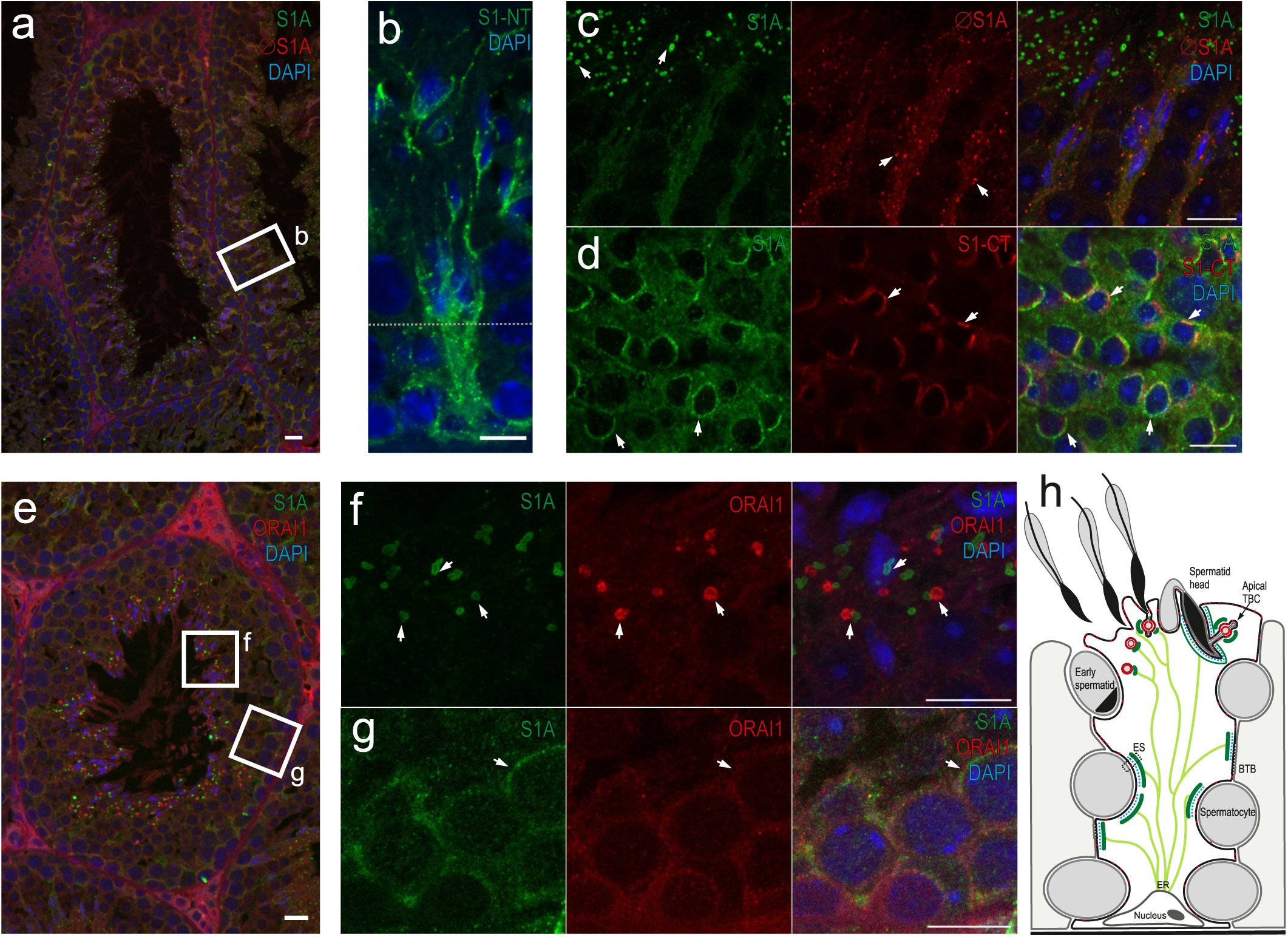
Stim1 variants detected in murine Sertoli cells by immunohistochemistry. (a) Overview of a cross-section through seminiferous tubules stained with *α*-Stim1A (S1A), *α*-ØStim1A (ØS1A) and DAPI. (b) Detailed display of a single Sertoli cell labelled with an N-terminal antibody against Stim1, lumen at top of image. (c) Detailed display of an apical region of [a] stained with *α*-S1A, *α*-ØS1A and DAPI. (d) detailed display of a basal region here stained with *α*-S1A, *α*-Stim1 (C-term) and DAPI. (e) overview of a cross-section through a seminiferous tubule stained with *α*-S1A, *α*-Orai1 and DAPI. (f) detailed display of an apical region of [e] (g) detailed display of a basal region of [e]. MIP, scale bars: 10 µm. (h) Schematic representation of membrane structures and distribution of domain A within a Sertoli cell. Adapted from “The endoplasmic reticulum, Ca^2+^ signaling and junction turnover in Sertoli cells,” with permission by Vogl et al. ^24^. Indicated in light green is the general ER as also seen in [b]. Dark green indicates specialized contact sites containing Stim variant (s) harboring domain A as observed in [c] and [d]. Shown in red is the distribution of Orai. The blue dotted lines symbolize actin filaments. ER: endoplasmic reticulum; ES: ectoplasmic specializations; BTB: blood testis barrier; TBC: tubulobulbar complex.

### STIM1A is a dominant-negative regulator of Orai– mediated Ca^2+^ entry and I_CRAC_

To analyze the impact of the inserted residues, we expressed either Stim1 or StimA in murine embryonic fibroblast (MEF) cells derived from *Stim1*^−/−^;*Stim2*^−/−^ mice (MEF S1/S2^−/−^) ^10^ and performed Fura-2 based Ca^2+^ imaging experiments. Figure 3a shows that re-expression of Stim1 in this background leads to recovery of the otherwise absent SOCE ^10^. In comparison to Stim1, StimA induces a reduced SOCE with a significantly decreased influx rate, peak and plateau (Fig. 3a,b), despite similar protein levels (Fig. 1g, see also below). We next co-expressed Stim1A while keeping Stim1 constant to investigate whether Stim1A is able to exert a dominant-negative effect on SOCE. Influx rate and peak indeed were significantly reduced, while the plateau reached similar levels to single Stim1 expression (Fig. 3a,b). To investigate whether Orai homologs are differentially affected by Stim1A, we expressed either *Stim* variant together with *Orai1, Orai2* or *Orai3* in HEK293 cells and to facilitate comparison, we quantified the resulting % change of the SOCE parameters. Overall, Stim1A led to a general decrease in SOCE parameters independent of the Orai homolog (Figs. 3c-f). For changes in absolute values, see Supplementary Figure S3. To confirm the dominant-negative nature of Stim1A, we performed siRNA experiments on cultured murine hippocampal astrocytes (Fig. 3g-i). SiRNA mediated downregulation (Fig. 3h) increased rate, peak and plateau of SOCE and verified the significance of Stim1A in primary cells. Subsequent analysis by whole-cell patch-clamp confirmed a significant decrease in Orai1 and Orai2 mediated current densities with Stim1A (Fig. 4a-f). The resulting IV relationships seen upon co-overexpression with Orai1 still displayed the typical inward rectification (Fig. 4c,f), suggesting no overt alteration of selectivity.

**Figure 3.**
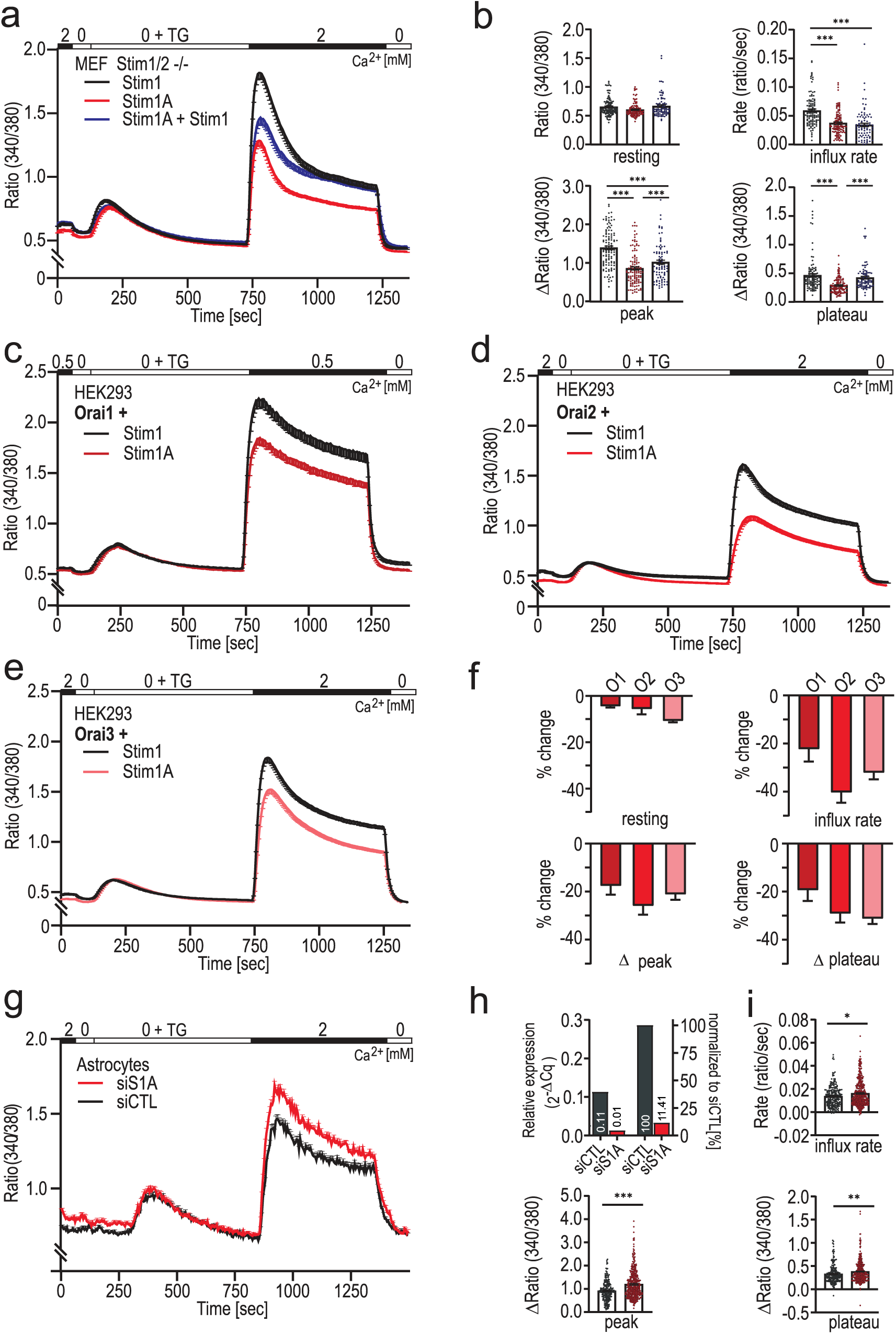
Stim1A reduces SOCE. (a) Traces showing average changes (mean±s.e.m.) in intracellular Ca^2+^ (Fura2 ratio) over time in response to perfusion of different [Ca^2+^]_o_ as indicated in the upper bar in MEF Stim1/Stim2 ^−/−^ cells transfected with *Stim1-* (black traces, n=111), *Stim1A* (red traces, n=116) or the combination *Stim1* with *Stim1A* (blue trace, n=86). (b) Quantification of changes in ratio of resting, influx rate (Δratio/time), Δpeak and Δplateau measured in a. (c-e) Traces showing average changes (mean±s.e.m.) in intracellular Ca^2+^ (Fura 2 ratio) in HEK293 cells co-transfected with either Stim1-(black traces, n=70-130) or Stim1A IRES-mCherry (red traces, n=66-90) and the indicated Orai1-3 homologs. (f) Relative change compared to the average of each respective control. (g) Traces showing average changes (mean±s.e.m.) in intracellular Ca^2+^ (Fura 2 ratio) in primary astrocytes after transfection with non-silencing or *exonA*-specific siRNA (siS1A). (h) Efficiency of siRNA as determined by qRTPCR. Quantification of parameters determined in (g). * p<0.05, ** p<0.01 *** p<0.001 Kruskal-Wallis Anova with Dunn’s multiple comparisons test (b,f) and Mann-Whitney test for (i). Data were obtained from three biological replicates with each three technical replicates and is shown as mean±s.e.m.

**Figure 4.**
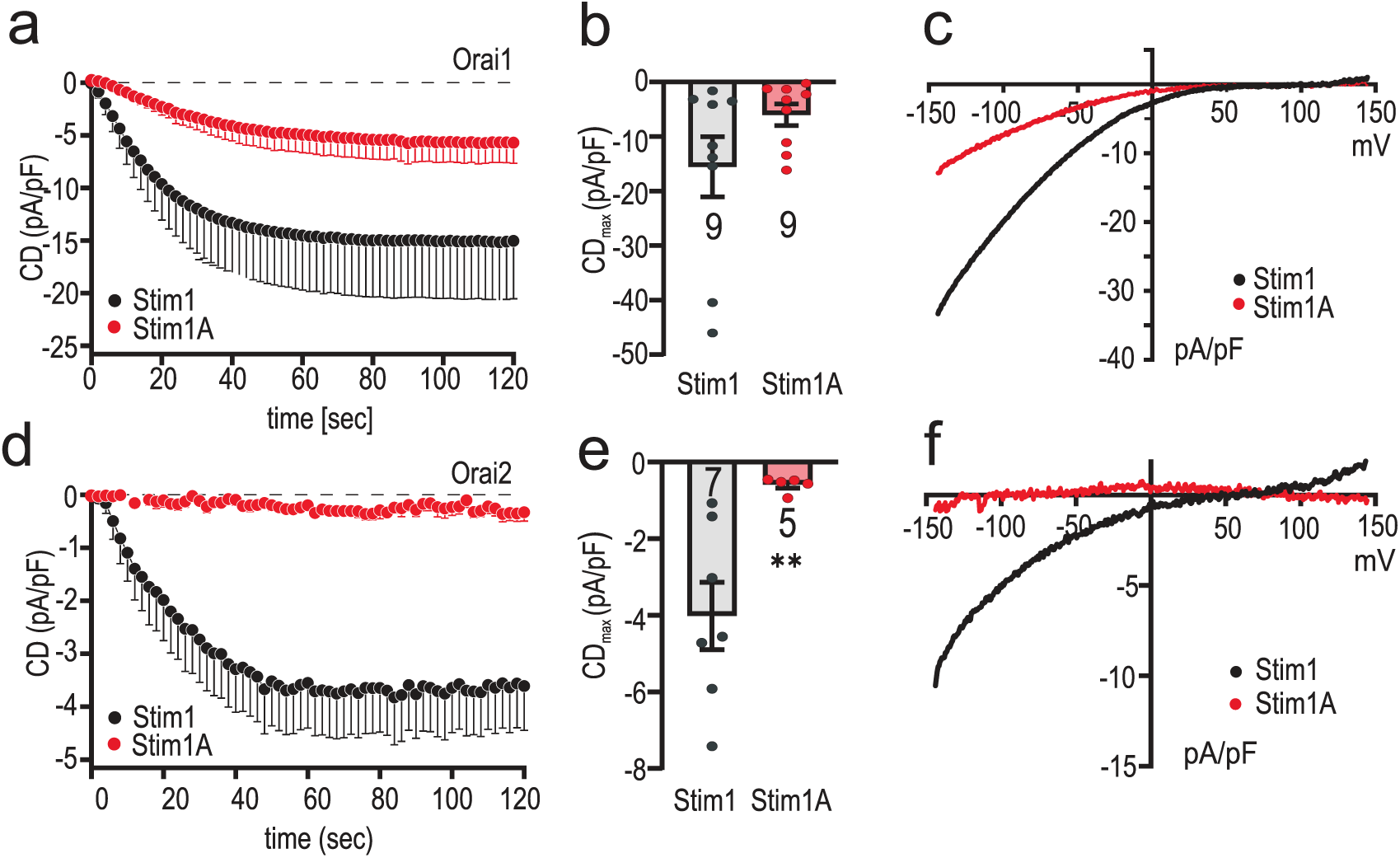
Inclusion of Domain A reduces I_CRAC_. (a) Average traces showing whole-cell current density (CD) over time extracted at −80mV in HEK293 cells co-transfected with Stim1 (black) or Stim1A (red) and Orai1. (b) Average maximum CDs recorded from cells measured in [a] (n within bars). (c) Current–voltage (I–V) relationship of all cells recorded in [a]. (d) Average traces showing whole-cell current density (CD) over time extracted at −80mV in HEK293 cells co-transfected with Stim1 (black) or Stim1A (red) and Orai2. (e) Average maximum CDs recorded from cells measured in [d]. (f) Current–voltage (I–V) relationship of all cells recorded in [a]. *P<0.05, **P<0.01 ***P<0.001, Student’s T-test. Data obtained are presented as mean±s.e.m

### STIM1A colocalizes with STIM1 in HEK cells and interacts with ORAI1

To investigate whether insertion of domain A leads to altered localization of STIM1, we transfected both variants into HEK cells lacking endogenous STIM proteins (HEK *S1/S2*^−/−^ cells ^18^) and investigated STIM1 – STIM1 *versus* STIM1A co-localization before and after store depletion. As depicted in Figure 5a, neither overt pre-clustering nor differential localization of STIM1A when compared to STIM1 was observed before store depletion. After stores were depleted, STIM1 or STIM1A clustered and co-localized to the same ER-PM junctions (Fig. 5b). Co-localization coefficients also revealed no differences (Fig. 5a,b). To investigate whether the reduced SOCE/I_CRAC_ is due to defective interaction or clustering with ORAI1, we analyzed colocalization and interaction with total internal reflection fluorescence microscopy (TIRFM). Both STIM1 and STIM1A co-localized with ORAI1 after store depletion at ER/PM junctions with no apparent difference in the Mander’s coefficients of co-localization (Fig. 5c,d). Quantification of FRET efficiencies (E-FRET) also indicated that interaction between the respective CAD (SOAR) domains of STIM1A and the C-terminal domain of ORAI1 was unaltered when compared to STIM1 (Fig. 5c,e).

**Figure 5.**
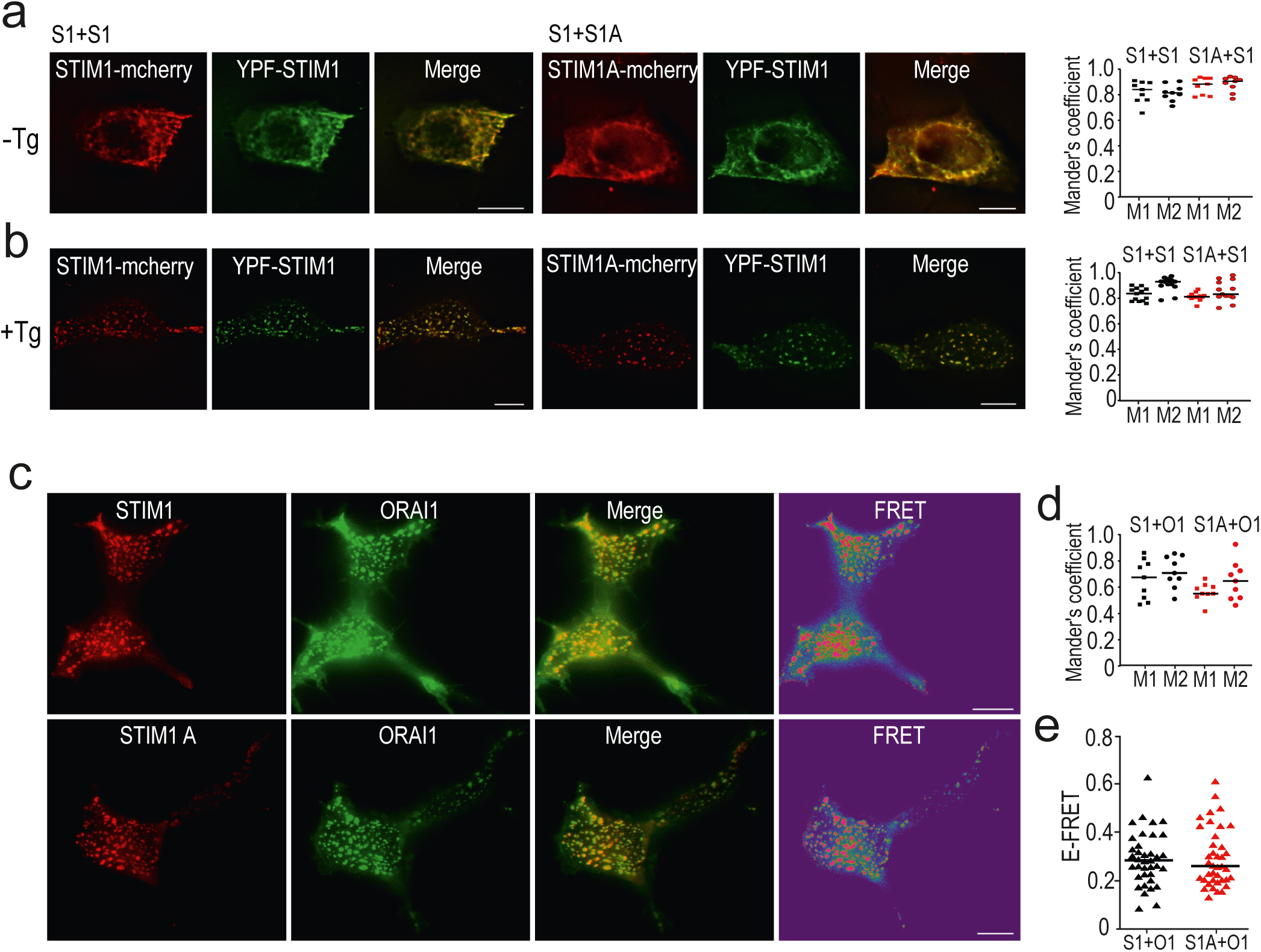
Co-localization and interaction analysis. Images showing representative HEK *S1/S2*^−/−^ cells co expressing YFP-STIM1 (green) or either STIM1-mCherry or STIM1A-mCherry (red) and merged images before (a) and after (b) stimulation with Tg. The rightmost panel show analyses of 9 cells using Mander’s overlap coefficients (M1 and M2) (-TG: S1+S1 M1=0,83 M2=0,87, S1+S1A M1=0,79 M2=0,89; +TG: S1+S1 M1=0,84M2=0,93, S1+S1A M1=0,81 M2=0,83). (c) Images showing representative HEK293 cells expressing Orai1-GFP (green = donor molecule) or either STIM1-mCherry (upper panel = acceptor molecule) or STIM1A-mCherry (lower panel = acceptor molecule) after stimulation with TG. Also shown is the overlay of the two channels, as well as the FRET signal. (d) Co-localization analysis of cells from c. For each condition 9 cells were analyzed using Mander’s overlap coefficients (M1 and M2) S1 M1=0,68 M2=0,71, S1A M1=0,55 M2=0,65. (e) Quantification of the FRET signal measured in c (STIM1: 0,28 n=37, STIM1A: 0,26 n=39). *P<0.05, **P<0.01 ***P<0.001 Mann Whitney test. Data was obtained from three independent transfections and is shown as single values with corresponding median. Scale bars, 10 µm.

### Specific residues in domain A mediate the reduced function

To further address the molecular mechanism of reduced SOCE and I_CRAC_, we asked whether the mere insertion of residues shortly downstream of the inhibitory domain (ID) is responsible for the phenotype or whether specific and conserved residues of domain A are required. As shown in Fig. 1f, domain A contains several highly conserved and also charged residues. The initial strategy included mutagenesis of the charged RRFSD stretch to alanine residues (AAAAA) or, alternatively, the downstream DDQS motif to alanine residues (AAAA) within the mCherry tagged construct utilized for the aforementioned FRET experiments (Fig. 6a; Fig. S5). The RRFSD/AAAAA mutant recovered the STIM1 wt phenotype, while mutation of DDQS/AAAA did not rescue the reduced SOCE (Fig. S5b,c,e). These results indicated that, indeed, specific residues and not a sole extension of the C-terminal domain encode the domain A phenotype. Further mutational analyses showed that the charged di-arginine motif (RR, with mutant QQFSD) was not responsible for reduced SOCE but rather that the FSD motif was a critical determinant of the STIM1A phenotype, as the FSD/AAA within STIM1A was sufficient to restore the STIM1 wt phenotype (Fig. S5b,e). Single point mutations were introduced within residues S502 and/or D503, both leading to enhancement of SOCE (Fig. S5d,f). As S502 potentially is modifiable by phosphorylation, a phospho-mimetic S502D mutant was also created but displayed a similar rescue as the S502A mutation, thus implying that phosphorylation of S502 is not the cause for the reduced function (Fig. S5d,f). The described functional screening was performed in HEK*S1/S2*^−/−^ cells with only endogenous ORAI channels mediating Ca^2+^ entry, thus leading to very small SOCE. We, therefore, confirmed the effects of STIM1A D503A in HEK cells stably over-expressing ORAI1 (HEKO1^29^). Here, we found a more pronounced difference between SOCE parameters of STIM1 and STIM1A, which were reversed with the D503A mutation (Fig. 6b). Concomitant measurement of mCherry fluorescence of the tagged constructs enabled control for equal protein expression (Fig. 6c) and quantitative analysis of all parameters confirmed the reversion of the STIM1A phenotype (Fig. 6d). In line with the Ca^2+^ imaging results, whole-cell patch-clamp analysis confirmed that STIM1A D503A also fully rescued the reduced current densities seen with STIM1A (Fig. 6e-g).

**Figure 6.**
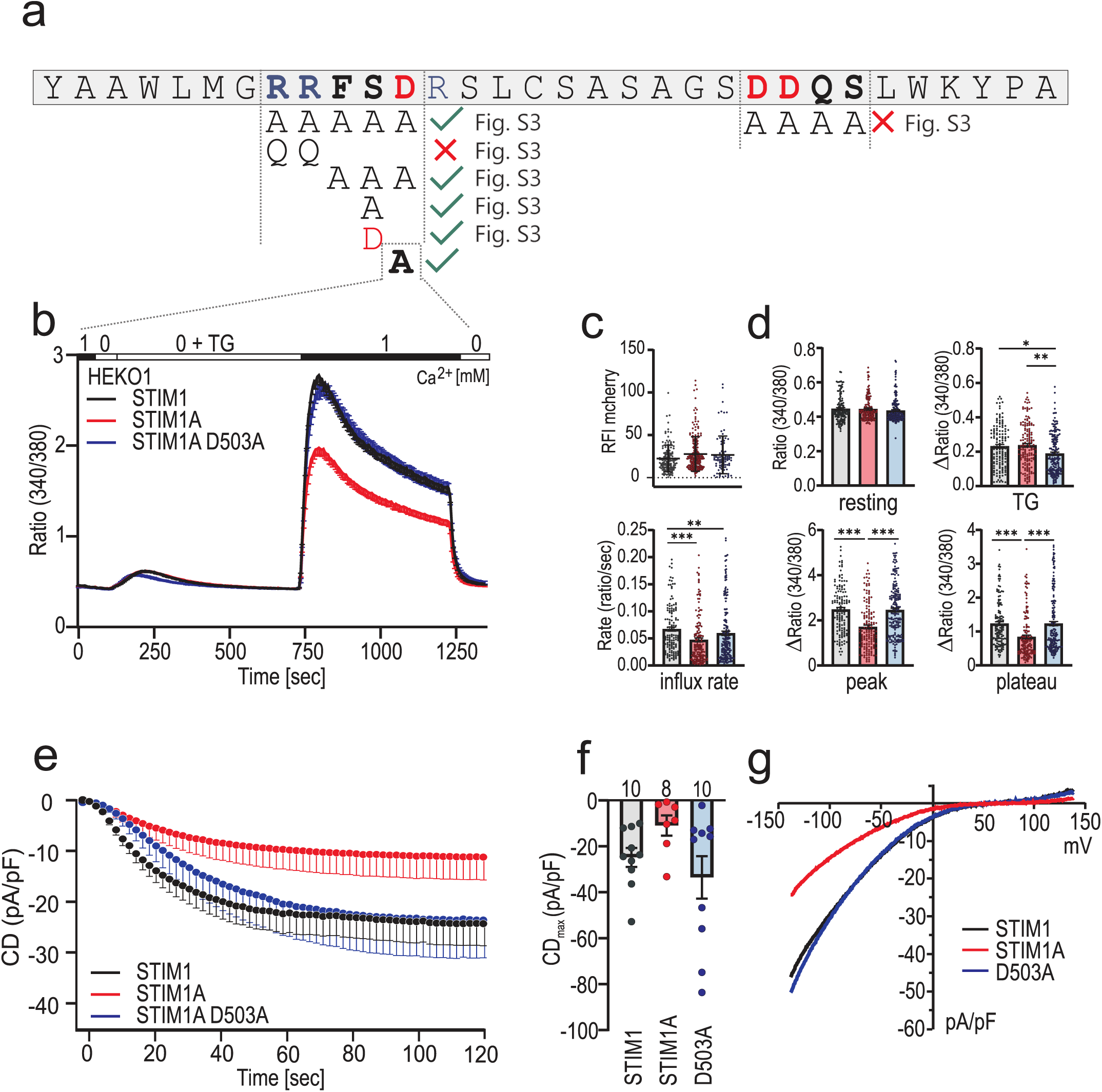
Mutation of D503 rescues phenotype of domain A. (a) Amino acids of exon A. Positive charged amino acids are marked blue, negative charged ones are shown in red. Mutated regions are in bold and different mutant combinations investigated in Fig. S5 are depicted below. (b) Traces showing average changes (mean±s.e.m.) in intracellular Ca^2+^ (Fura2 ratio) over time in response to perfusion of different [Ca^2+^]_o_ as indicated in the upper bar with constructs as indicated expressed in HEKO1. STIM1 (black, n=145), STIM1A (red, n=157) and STIM1A D503A (n=185). (c) Relative fluorescence intensities of mCherry-tagged constructs measured in (b). (d) Quantification of changes in ratio or ratio/time measured in b. (e) Average traces showing whole-cell current density (CD) over time extracted at −80mV in HEKO1 cells transfected with STIM1 (black), STIM1A (red) or STIM1A D503A (blue). (f) Average maximum CDs recorded from cells measured in e (n within bars). (g) Current–voltage (I– V) relationship of all cells recorded in e. *P<0.05, **P<0.01 ***P<0.001 Kruskal-Wallis Anova with Dunn’s multiple comparisons test.

### Domain A interferes with full gating of ORAI1

Gating of ORAI1 by STIM1 is mainly governed by a primary interaction of the STIM1 CAD/SOAR domain with the C-terminus of ORAI1 and potentially directly with the “nexus” site (amino acids 261-265 of ORAI1^30^). This site elicits a conformational transduction pathway via trans-membrane domain TM4→TM3→TM1 helix to open the pore (reviewed recently in ^4,31^). However, several groups have suggested that the CAD domain also interacts with the **E**xtended **T**ransmembrane **O**rai1 **N**-terminal (ETON) region (aa 73–90) [^32–34^] and that this region is critical for gating as well as fast Ca^2+^ dependent inactivation. ORAI1 gating may thus be stabilized by a STIM1-meditated bridging between the cytosolic TM1 and TM4 extended helices of ORAI1, thereby applying a force at the helical TM1 extension to stabilize the open pore state ^32,35^ or alternatively between TM1 and the TM2-3 linker (see discussion). Domain A is inserted eight residues downstream of an acidic inhibitory domain (ID), which has been also been postulated to functionally interact with conserved residues (W76;R77;K78) of the extended ORAI1 pore helix (TM1, ETON region) to mediate fast Ca^2+^ dependent inactivation (FCDI) ^36^. We therefore asked if domain A would interfere with a stabilizing interaction between upstream STIM1 residues and the ORAI1 ETON region. If this were the case, an N-terminal gate-modified ORAI1 such as ORAI1 R77E might mask the STIM1A phenotype. Using Ca^2+^ imaging, we observed a 35–40% reduced STIM1 mediated SOCE with ORAI1 R77E (Fig. 7a,d). This difference indeed was absent when STIM1A was co-expressed with ORAI1 R77E (Fig. 7b,d), but could be restored with the single D503A mutation within domain A (Fig. 7c,d). Patch-clamp analysis in HEK*S1/S2*^−/−^ cells confirmed the reduced currents with STIM1A (human) (Fig. 7e,g) similar to those observed with Stim1A (murine) in HEK*wt* cells (Fig. 4). Co-expression with ORAI1 R77E leads to a strong current reduction compared to ORAI1 (Fig. 7e,g and ^33^). However, STIM1A does not lead to a further reduction in the background of R77E (Fig. 7f,g). These results suggest that insertion of domain A affects the interaction of STIM1 CAD residues and R77 of ORAI1, thus either reducing or destabilizing full gating of ORAI1.

**Figure 7.**
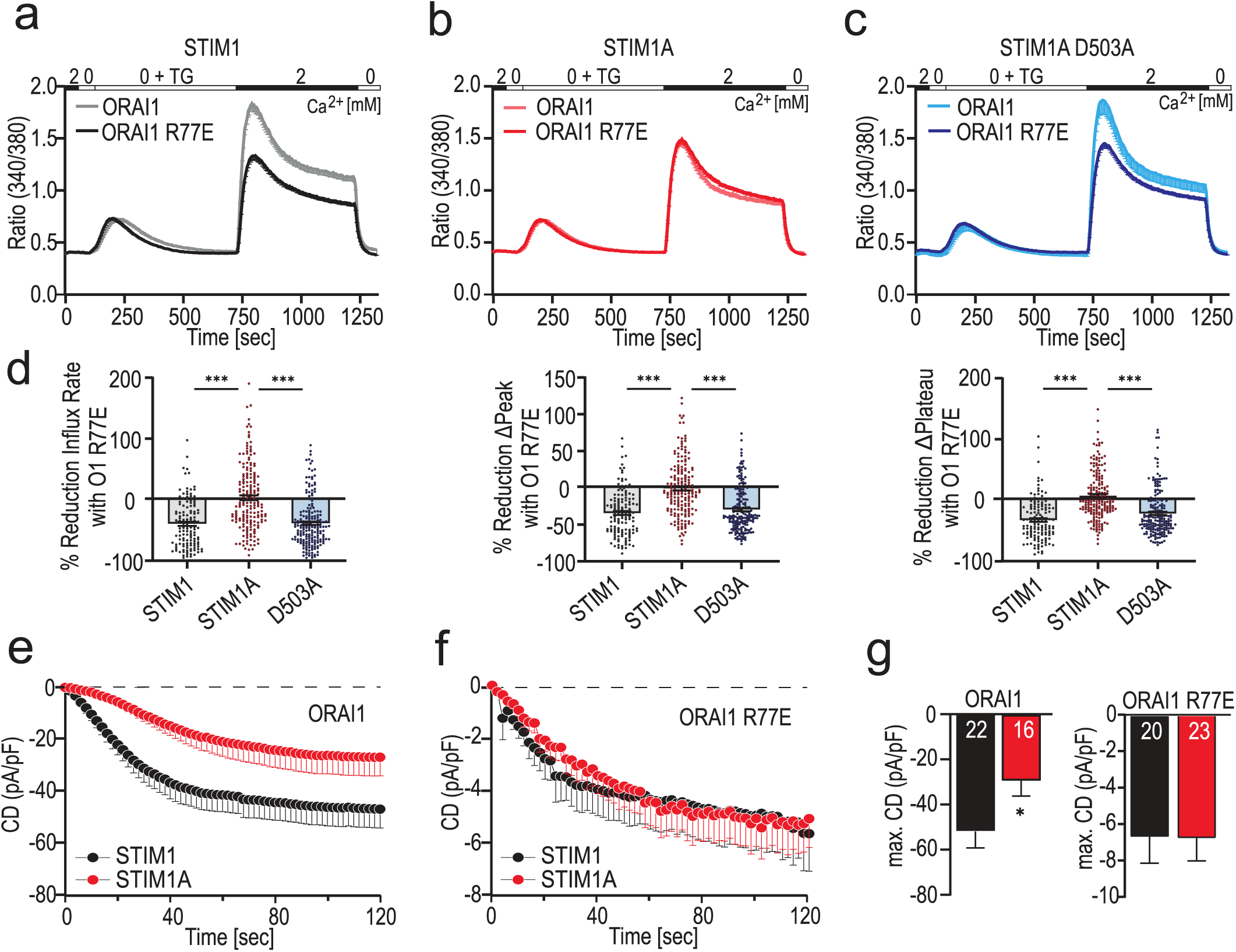
Mutation within the ORAI1 ETON region mask the STIM1A phenotype. (a-c) Traces showing average changes (mean±s.e.m.) in intracellular Ca^2+^ (Fura2 ratio) over time in response to perfusion of different [Ca^2+^]_o_ as indicated in the upper bar with constructs as indicated expressed in HEKS1/S2^−/−^. (d) Quantification of the relative reduction of ratio/time or ratio for cells (135<n<202) measured in a-c. (e,f) Average traces showing whole-cell current density (CD) over time extracted at −80mV in HEKS1/S2^−/−^ cells co-transfected either with STIM1 (black) or STIM1A (red) and ORAI1 (e) or ORAI1 R77E (f) and recorded using extracellular solution containing 2 mM Ca^2+^. (g) Average maximum CDs recorded from cells measured in (e,f) (n within bars). *P<0.05, **P<0.01 ***P<0.001. d: Kruskal-Wallis Anova with Dunn’s multiple comparisons test, in g: unpaired T test with Welch’s correction.

## Discussion

In our study we report the identification and functional characterization of a novel STIM1 splice variant, STIM1A, where 31 amino acids are inserted into the cytosolic C-terminal domain downstream of the CAD/SOAR domain and immediately downstream of the highly charged acidic inhibitory domain (ID) domain. We show that while *exon A* is expressed in many tissues, its relative expression varies among tissues and cell types and is particularly high in testes, heart, astrocytes and kidney. Within the genomic structure of both the murine and human *STIM1* gene several additional splice variants are predicted with both alternative and mutually exclusive splice events being present. The identification of several new exons necessitates a new numbering nomenclature of the twelve exons of the conventional Stim1. Although alternative *5’UTR* regions can be spliced in before exon 2 and lead to translation of an internal methionine, no new exon appears to be spliced in within the N-terminal region. Splice-insertion of *exon A* within full length *Stim1* translates to a protein detected within testes (Fig. 1h). Testes also express splice combinations leading to N-terminally truncated Stim1 and Stim1A, although at low levels as detected by qualitative and quantitative RT-PCR (Fig. S1c,d). While the relative amount of N-terminal spliced variants in several investigated tissues is low; this variant will require further investigation. Functionally, Stim1A reduces SOCE and I_CRAC_ in a sequence specific dominant-negative manner, with the conserved FSD motif within domain A being essential and either S502A or D503A point mutants sufficient for phenotype reversion. We found no overt defect in FRET between STIM1A and ORAI1 after store depletion in HEK cells. Furthermore, STIM1A did not lead to a reduction in SOCE or I_CRAC_ in conjunction with the ORAI1 R77E mutation. One possible explanation is that domain A prevents a stabilizing interaction between the ETON region and CAD/SOAR, necessary for full and stabilized gating. The finding that normal function is restored with mutation of the FSD motif or with single residue mutations (S502A or D503A) could be due to a structural change resulting in regained accessibility of the interacting upstream residue(s) (model Fig. S6). An alternative explanation of the STIM1A phenotype includes recruitment of an additional interaction partner(s) which binds to domain A and reduces SOCE. Mutations of FSD would then disable this interaction. However, the fact that with ORAI1 R77E the phenotype is masked argues against a binding partner responsible for the heterologous SOCE phenotype.

Stim1 is known to play a critical role in both in astrocytic Ca^2+^ signaling and gliotransmitter release ^37^ as well as in testicular cord formation in mouse testis as knock-down of total Stim1 caused defects in gonadal development, loss of interstitium, failed angiogenesis and increased levels of reactive oxygen species ^26^. Which of the above mentioned functions seen upon loss or downregulation of total Stim1 is due loss of Stim1A is unknown and requires further investigation. Remodeling of intracellular junctions within the seminiferous epithelium is critical for spermatogenesis as there is a constant need for disassembly and reassembly of junctional complexes between neighboring Sertoli cells as germ cells are translocated from basal to adluminal compartments, but little is known about the molecular mechanisms driving this process. During spermiation, large adhesion junctions between Sertoli cells and late spermatids need to be internalized ^38^. The intercellular adhesion with differentiating germ cells and between Sertoli cells are named “ectoplasmic specializations” (ES) and are defined as a link of ER cisternae to actin filaments, which in turn are linked to regions of the Sertoli cell plasma membrane. The mostly apically localized so-called “tubulobulbar complexes” (TBCs) are specialized ER regions internalizing intercellular junctions after sperm release ^27^ and have some molecular and structural similarities to podosomes of other cells ^39^ as well as sharing common features with clathrin-mediated endocytosis ^40,41^. They also contain specialized membrane contact sites (MCS) devoid of actin which contain essential proteins of the lipid transfer machinery ^42^ (for schematic visualization see Fig. 2h). Using a combination of different validated Stim1 antibodies as well as a domain A specific antibody, we find a highly specialized targeting of Stim1 splice variants containing domain A to different specialized ER regions (Fig. 2). All antibodies against Stim1 recognize general ER structures within Sertoli cells, while only Domain A as well as the monoclonal C-terminally directed STIM1 antibody highlight ES structures adjacent to germ cell nuclei. These structures are not strongly labelled with the ØA-antibody, suggesting that Stim1A with an intact C-terminus (but not Stim1wt) targets to the ES (Fig. 2). Within the TBCs, close to spermatids with condensed chromatin, we find dense packing of domain A within ER structures adjacent to tubules of plasma membrane highlighted by the Orai1 antibody. These localized ER structures also express early endosome markers such as Rab5 and EEA1^43^, but are not recognized by the N-terminal STIM1 antibodies; therefore, these domains likely contain the alt1-Stim1A splice combination. Whether a permanent association with these Stim1 variants and Orai in the PM is required for endocytosis and is related to the finding that Orai channels are critical for receptor-mediated endocytosis of albumin in also in proximal tubular epithelial cells ^44^ requires further investigation and generation of splice-specific knock-out models. What also remains unclear is how the dramatic remodeling and various assemblies at these specializations are kept in a dynamic equilibrium and how they are regulated. Wales et al. have proposed a mechanism termed Ca^2+^-mediated actin reset (CaAR) to transiently immobilize organelles and to drive reorganization of actin ^45^. It is possible that for some membrane contact sites, Stim1A with reduced Ca^2+^ entry is required, while for TBC regulated internalization, a disabled EF-hand leading to strong Orai1 coupling may drive membrane invagination in TBC regions lacking actin (Fig. 2h). Within the native environment of a Sertoli cell, Stim1 splicing therefore (1) differentially regulates SOCE to most likely fulfill specific contact site Ca^2+^ requirements and (2) adds zip-codes to specifically target Stim1 variants to different membrane contact sites.

## ACKNOWLEDGEMENTS

We thank Phillip Knapp, Pauline Schepsky, Drs. Olga Ratai, Jens Rettig and Jutta Engel for help with the Cell Observer and LSM microscopes, Zoe Schmal and Dr. Claudia Rübe for help with testes samples, Alina Gilson and Maik Konrad for initial experiments, Lukas Jarzembowski for help with formatting the manuscript. We also thank Dr. Wayne Vogl and members of the Hoth’ and Niemeyer’ labs for critical input, reading of the manuscript and discussions.

## AUTHOR CONTRIBUTIONS

BAN and MLK designed experiments and wrote the manuscript with help from DA and AL. MLK and KF performed imaging, PCR, immunhistochemical and biochemical experiments, YS provided astrocytic cultures, MJ produced antibodies, DA and AL performed patch-clamp experiments. All authors conducted data analysis.

## COMPETING INTERESTS

The authors declare no competing or financial interests.

## FUNDING

The research was funded by the DFG SFB894 (A2) and SFB1027 (C4, C7) to BAN. Mona L. Knapp (née Schoeppe) received support by IRTG1830 and SFB894.

## Material and Methods

### Cell lines and Transfection

All cell lines were cultivated at 37 °C, 5%CO_2_ in a humidified incubator in respective medium (HEK293 minimum essential medium, HEK293*STIM1/2*^−/−^ dulbecco’s modified eagle’s medium, MEF*Stim1/2*^−/−^ Dulbecco’s modified eagle’s medium, HEK O1 minimum essential medium. All culture media were supplemented with 10% fetal calf serum. For passaging cells were detached using trypsin. For transfections HEK cells were transfected via electroporation unsing the Amaxa® Nucleofector II® (Lonza) according to the user manual. Amount of DNA used was 1 µg/1.000.000cells. MEF cells were transfected using jetPRIME® (Polyplus transfections) transfection reagent following the manufacturer’s protocol. siRNA (#1+#2, 20 nM total) or siCTL (20nM) were transfected 24 h before recording using Interferin® (Polyplus transfections). Cells were analyzed 20–24 h after transfection. DNA constructs, primers and siRNA are listed in Supplementary Table 1.

### Hippocampal astrocyte cultures

Hippocampal astrocytes were prepared from from C57BL/6 mice of either sex at postnatal days P0 to P1. Hippocampi were dispersed using a cell strainer in DMEM (Invitrogen). After centrifugation for 10 min at 1900 rpm at RT cells were resuspended and plated in collagen (0,5 mg*/*mL; Bioscience) coated culture flasks and cultured at 37 °C in culture medium (DMEM + 10% FCS + 0,1% P/S + 0,1% Mito Serum Extender) at 8% CO_2_ until cultures became confluent (5-7 DIC). Cells were subsequently trypsinized, plated on collagen-coated coverslips and grown until confluent (5-7 DIC).

### PCR and Quantitative real-time PCR

For cDNA synthesis cells were first harvested in TRIzol (Life technologies) and RNA was isolated according the manufacturer’s instructions. The cDNA transcription was obtained using SuperScriptTMII Reverse Transcriptase (Life technologies). For analytic PCR DreamTaq Green PCR Master Mix (Thermo Fisher Scientific) was used. Amplifications were analyzed in GTQ-agarose gels with Safe-Red DNA dye. qRT-PCRs were performed using QuantiTect SYBR Green Kit (Qiagen) and a CFX96 Real-Time System (Bio-Rad). All Primers are listed in Supplementary Table 3. Using the ΔCq (quantification cycle) method results were normalized to TBP, whereas normalization with HPRT1 showed comparable findings. Values are shown as 2^−(ΔCq)^.

### Western Blot/antibodies

Transfected cells were washed with PBS, detached with a cell scraper and transfered in ice cold RIPA Buffer containing 150 mMNaCl, 50 mMceTris-HCl, 1% Nonidet P 40, 1% Trition and protease inhibitor cOmplete(tm) (Sigma-aldrich). For lysis the cells were frozen 10 min at −80 °C and vortexted (3x 20 s). Murine tissues were washed in PBS and lysed using a micro homogenizer (neoLab). Lysed cells were spin down at 12,000 RPM (4 °C) for 30 min. Proteins were linearized in Tris/Glycine Bufferat 95 °C for 5 min, electrophoresed in 7% SDS/PAGE, and blotted onto a nitrocellulose membrane (GE Healthcare). Membranes were blocked with 5% skimmed milk in TBST Buffer. Primary and secondary antibodies used for Western blot are listed in Supplementary Table 2.

### Fluorescent based Ca^2+^ Imaging

Cells were loaded with 1 µM Fura2-AM for 30 min at RT. For perfusion a fully automated perfusion system (ALA Scientific Instruments) was used. Cells were flushed with 1mL of Ca^2+^ Ringer of different concentrations. The Ca^2+^ Ringer solution contained (in mM): 155 NaCl, 2 MgCl_2_, 10 glucose, 5 Hepes and 0.5-2 CaCl_2_ (= 0.5-2 Ca^2+^ Ringer) or no CaCl_2_, but 1 EGTA plus 3 MgCl_2_ instead (=0 Ca^2+^ Ringer) (pH 7.4 with NaOH). To block the SERCA 1 µM TG in 0 Ca^2+^ Ringer was used. Images were taken at 340 and 380nm every 5 s at RT and analyzed with VisiView® Software. To quantify 340/380nm signal ratio IgorPro was used. Parameters analyzed were the average basal signal, the maximal Tg-induced peak, maximal and plateau after readdition of 0,5–2mM [Ca^2+^]_o_ and influx rate. The minimal signal before addition of Tg or 0,5mM [Ca^2+^]_o_ was subtracted respectively from TG Peak or Ca Peak/Plateau to determine the Δs.

### TIRF microscopy for FRET analysis

HEK293 cells were transfected with 3 µg STIM1-mCherry-pmax (containing mCherry after position L599^46^) or STIM1A-mCherry-pmax (acceptor) and 1 µg Orai1-GFP-pmax (donor) 24 h before measurements. Prior to the measurements TG sensitive stores were depleted using 1 µM TG in 0 [Ca^2+^]_o_ Ringer. For recording fluorescence images the Leica AM TIRF MC system was used. Imaged were taken with a 100 × 1.47 oil HCX PlanApo objective. The GFP-Donor signal was used to determine the TIRF focal plane. For each cell three imaged were taken: I. GFP excited at 488nm(suppression filter BP 525/50) II. Mcherry excitation wavelength at 561nm (suppression filter BP 600/40) III. FRET excitation with a 488nm laser and suppression filter BP 600/40. To calibrate laser intensities and excitation durations for each day of experiments single transfected cells were used, only expressing the donor or acceptor construct. Calibrated Parameters were constant for all channels and images taken. For image acquisition and analysis the LAS (Leica Application suite) FRET module was used. To calculate FRET efficiency according to van Rheenen et al ^47^. All images were corrected for background signal, bleed through and crosstalk factors.

### Live cell imaging

HEK STIM1/2 ^−/−^ cells were transfected with SP-YFP-*STIM1* and *STIM1*-mCherry or *STIM1A*-mCherry. Fluorescence was detected using the wide-field epifluorescence microscope cell observer A1 (Zeiss) with the Fluar 40×/1.3 M27 oil objective. To detect YFP fluorescence filtercube 54HE and LED 470 (470/40) was used, filtercube 56HE and LED N-White + Ex (556/20) for mCherry. Images were taken as Z-stacks, and processed as a maximum intensity projection (MIP) for visualization. Colocalization was analyzed using the Fiji plugin JACoP form a single stack. Threshold was set as mean plus three times SD.

### Immunocyto-, Immunohistochemistry and antibodies

All steps were executed with pre-chilled solutions and on ice. Cells seeded on glass coverslips were fixated with MeOH and 5% acetic acid for 10 min at −20 °C. Paraffin embedded tissues were cut into 4 µm thick sections. Sections were dewaxed using xylene and hydrated in descending ethanol series. To de-mask antigens, sections were incubated in citrate buffer at 95 °C for 1 h. Un-specific epitopes of cells as well as tissue sections were blocked using 4% BSA in PBS for 30 min at room temperature. Primary antibodies and secondary antibodies were diluted in blocking solution and incubated over night at 4 °C (primary antibody) and for 1 h at room temperature (secondary antibody). Cells and tissue sections were mounted with ProLong™ Gold Antifade Mountant with DAPI (Thermo Fisher Scientific). Primary and secondary antibodies used for immunochemistry are listed in Supplementary Table 2. Images were acquired using the Zeiss cell observer A1 as described above (GFP: filtercube 38HE, LED 470 (470/40) or using a confocal microscope LSM 710 (Zeiss) with a ×20/1 numerical aperture (NA) and a ×63/1.4 NA oil objective and processed using Fiji ^48^. Images were taken as z-stacks with an optical slice thickness of 0,32 µm (LSM) and are shown as MIPs.

### Patch Clamp electrophysiology

Recordings were performed at room temperature in the tight-seal whole-cell configuration as in Kilch et al. ^29^. Briefly, the indicated cells were transfected as above and recordings were done 24 h later with an EPC-10 patch-clamp amplifier controlled by Patchmaster software (HEKA). Series resistance was compensated to 85% for the transfected HEK293 cells. Immediately after establishing whole-cell configuration, linear voltage ramps from −150mV to 150mV (50 ms duration) were applied every 2 s from a holding potential of 0mV for the indicated time period. Currents were analyzed at − 80mV and 120 s after break-in. The pipette solution contained the following (in mM): 120 Cs-glutamate, 3 MgCl_2_, 20 Cs-BAPTA, 10 Hepes and 0.05 IP_3_ (pH 7.2 with CsOH). Bath solution contained (in mM): 120 NaCl, 10 TEA-Cl, 2 CaCl_2_ (or 10 CaCl_2_), 2 MgCl_2_, 10 Hepes and glucose (pH 7.2 with NaOH).

## Supplementary Figures and Tables

**Supplementary Figure 1.**
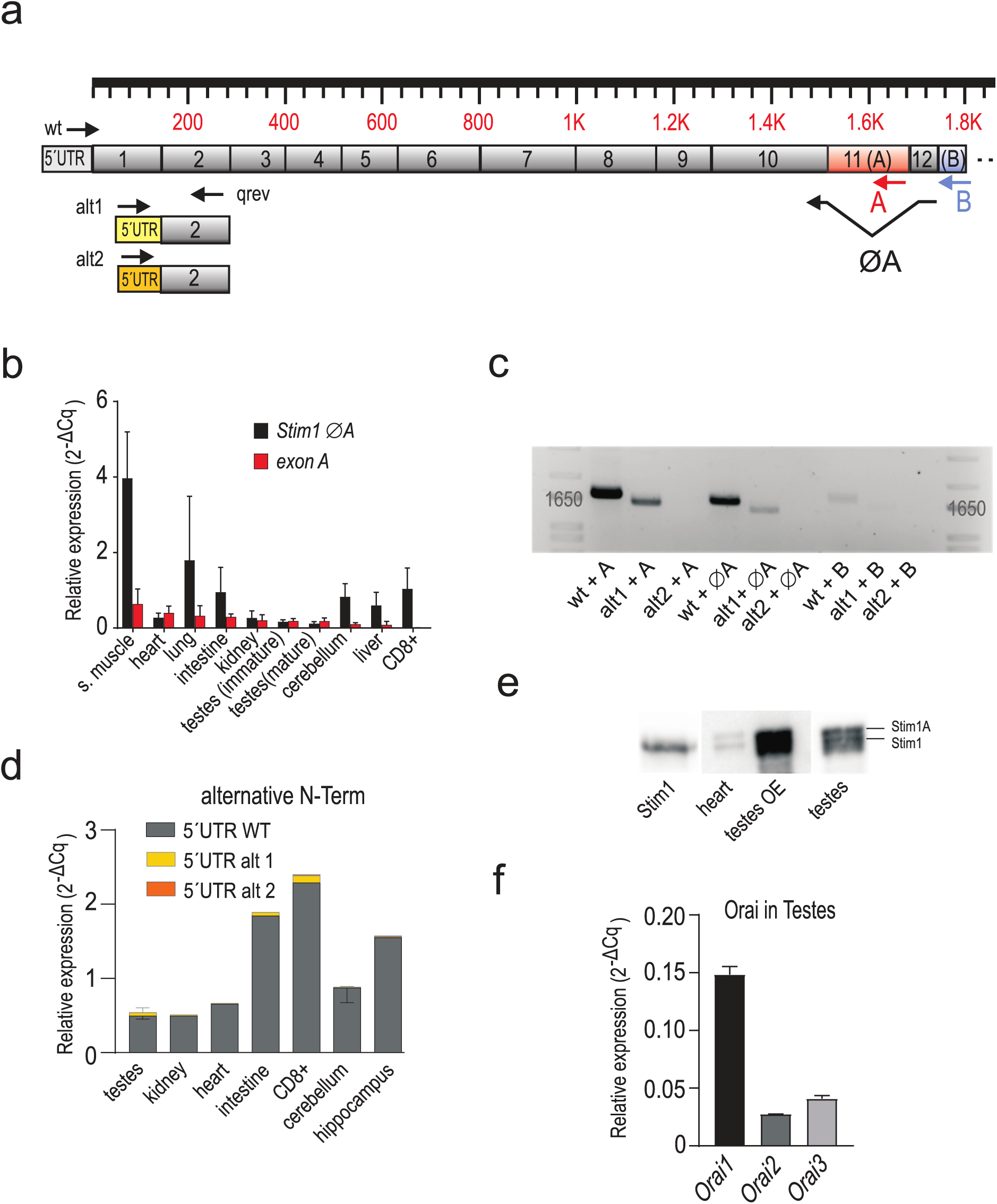
PCR analysis of Stim1 and domain A. (a) Schematic exon structure of *Stim1* with conventional exons depicted in grey. Highlighted in red is the alternative exon 11 (A) and the alternative exon 13 (B). Alternative usage of 5’UTR regions are indicated with alt1 and alt2. Annealing sites of primers are depicted as arrows. (b) Relative expression of *exon A* and *ØA* (± SD) in tissues and cells using cDNA’s derived from 3-4 mice normalized to TBP (c) Analytic PCR with reverse primers flanking the A splice site or within exon A or exon B in combination with forward primers specific for *5’URT wt, alt1* or *alt2* as also depicted in (a) using cDNA of murine testis. (d) Quantification of alternative N-term splicing via qRT-PCR. (e) Western blot analysis of lysates from HEK STIM1/2 ^−/−^ cells overexpressing Stim1 as well as whole tissue lysates of heart and testes showing bands for Stim1A as well as Stim1. The middle part of the blot shows testis as overexposed (testes OE) and was repeated with lower exposure time (right part of the blot). (f) Expression of the Orai homologs in murine testes. Expression levels were normalized to TBP.

**Supplementary Figure 2.**
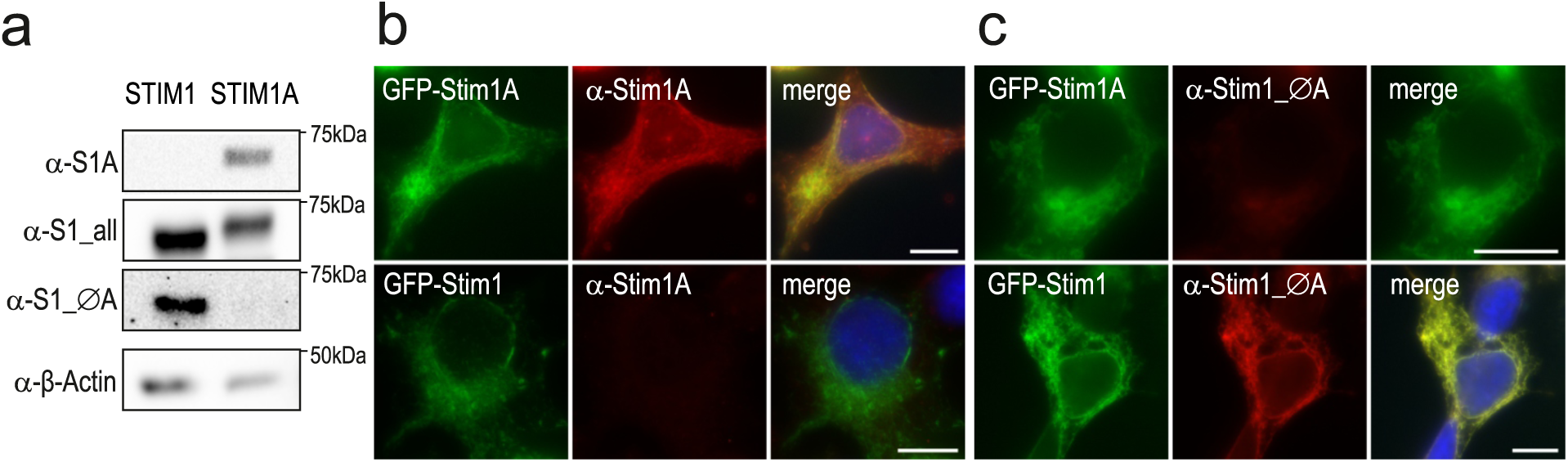
Test of the A specific and ØA Antibody. (a) Western blot showing Stim1 and Stim1A after heterologous expression in HEK293. Blot was probed with a domain A specific antibody, a ØA antibody and a Stim antibody binding its very N-term. (b) Immunocytochemistry of HEK S1/S2 ^−/−^ transfected with GFP-STIM1 or GFP-STIM1A after staining with a domain A specific antibody. (c) Immunocytochemistry of HEK S1/S2^−/−^ transfected with GFP-STIM1 or GFP-STIM1A after staining with a ØA antibody. MIP, Scale bars: 10 µm.

**Supplementary Figure 3.**
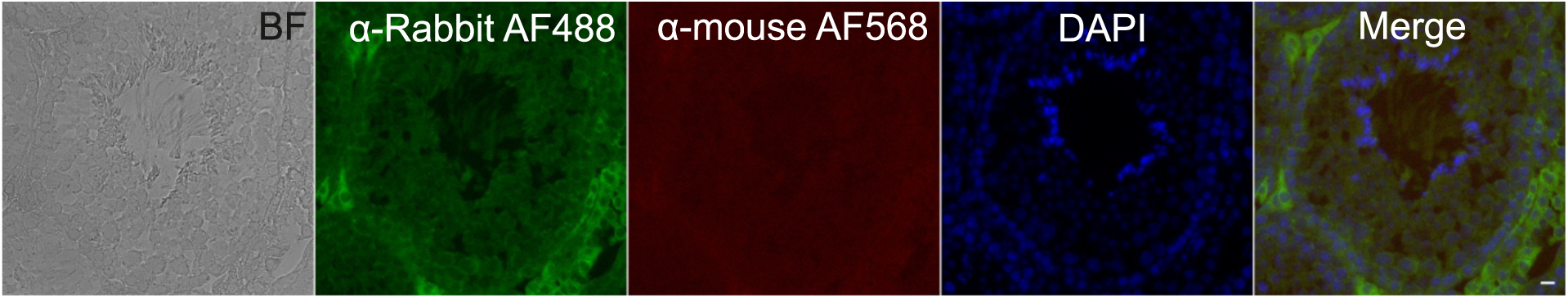
Immunohistochemical non primary antibody control. Murine testes were stained with only the secondary antibody goat *α*-rabbit AF488, goat *α*-mouse AF568 (Invitrogen) and DAPI. Single stack, Scale bars, 10 µm.

**Supplementary Figure 4.**
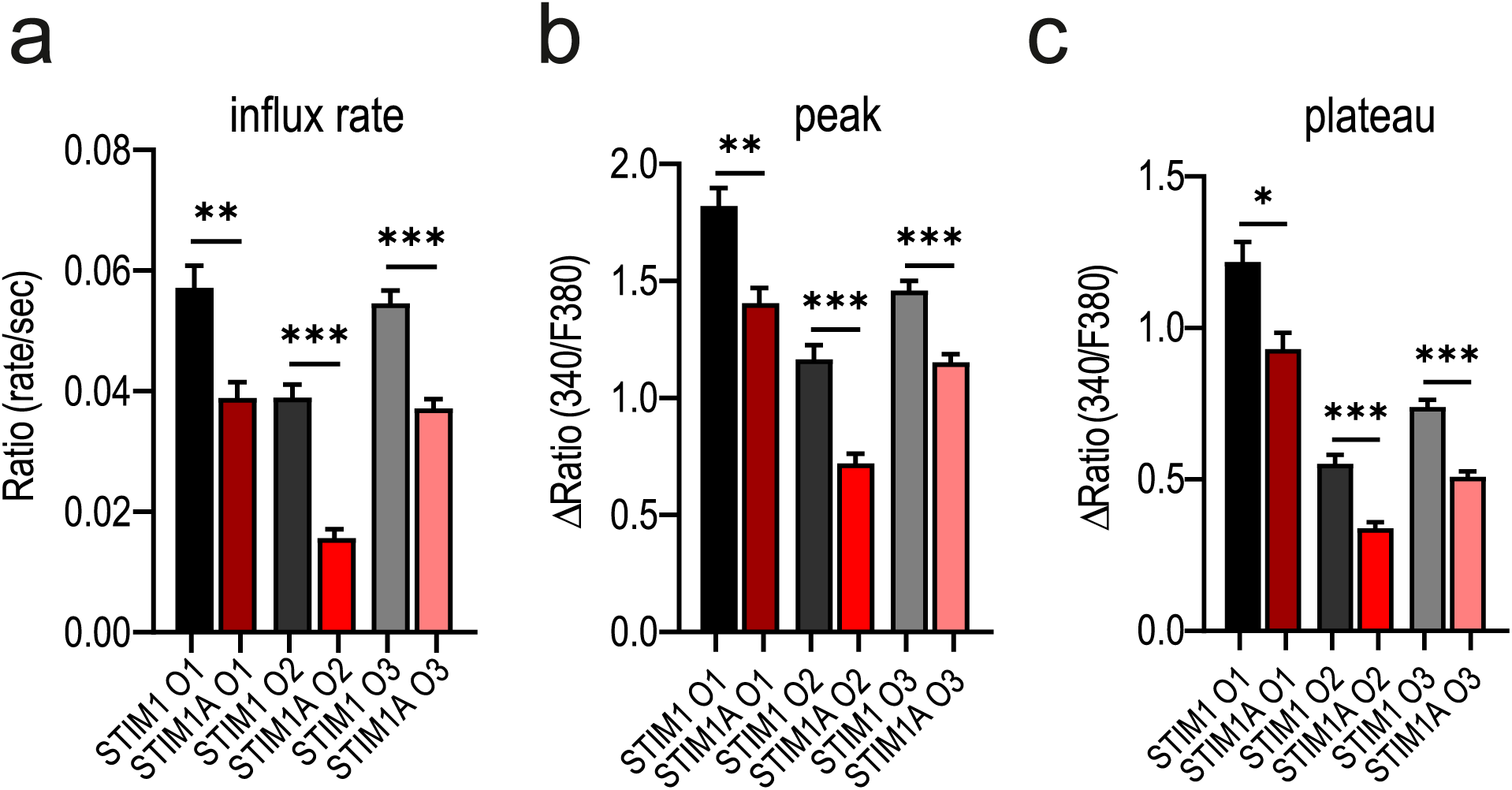
Stim1A reduces SOCE independent of the Orai homolog. Quantification of changes in ratio or ratio/time for the SOCE parameters ((a)influx rate, (b) Peak, (c) Plateau) measured in Fig. 3 c-e. * p<0.05,** p<0.01 *** p<0.001, Mann Whitney test for pair-wise comparisons with each Orai homolog. Data was obtained from three technical from three biological replicates and is shown as mean±s.e.m.

**Supplementary Figure 5.**
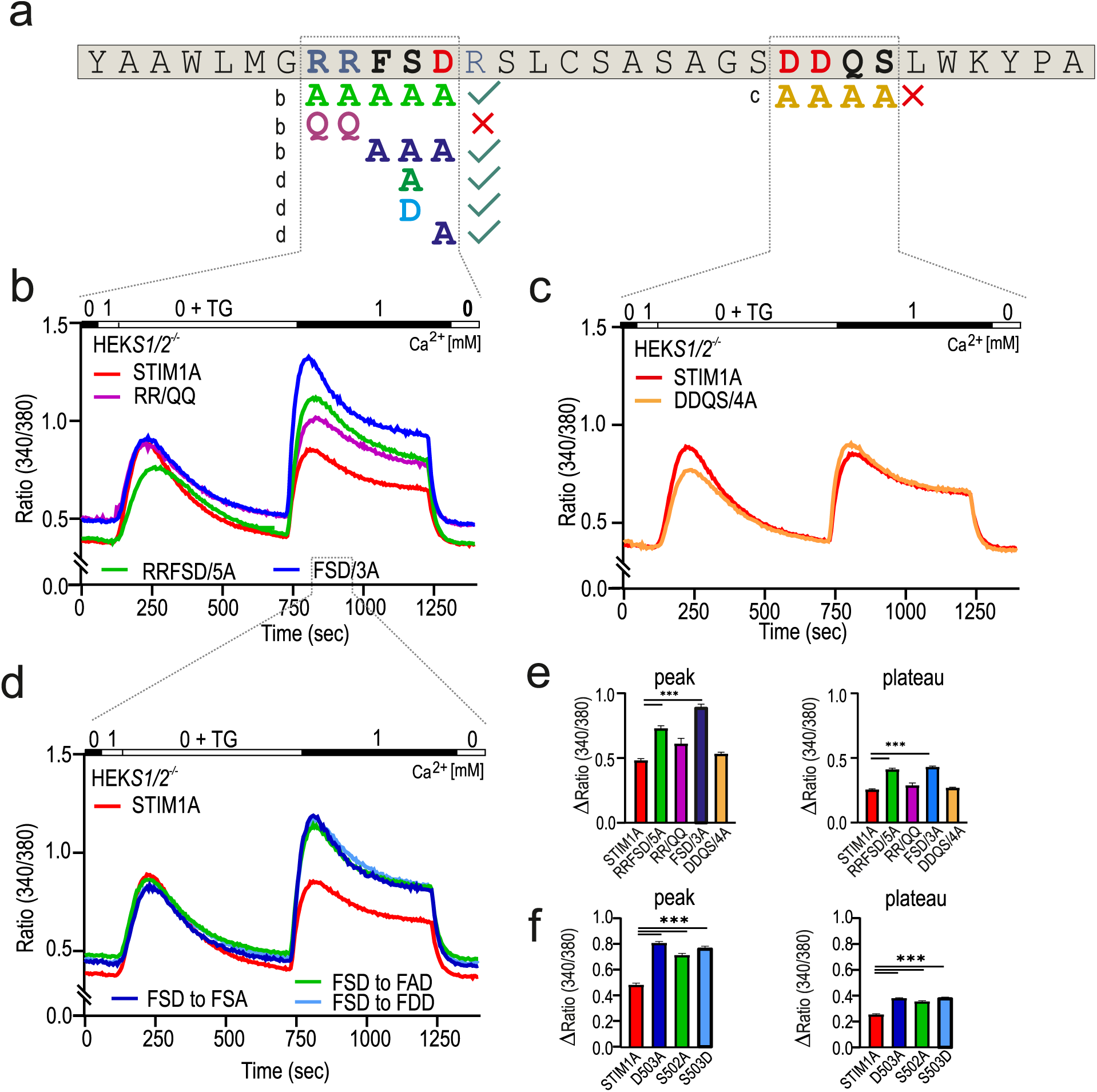
Site-directed mutagenesis of exon A. (a) Amino acids of exon A. Positive charged amino acids are marked blue, negative charged ones are shown in red. Mutated amino acids depicted below are marked in the color code used for b-f. (b-d) Traces showing average changes (mean±s.e.m.) in intracellular Ca^2+^ (Fura2 ratio) over time in response to perfusion of different [Ca^2+^]o as indicated in the upper bar with mutant constructs as indicated expressed in HEK*STIM1/STIM2*^−/−^ cells. (e) Quantification of changes in ratio measured in b and c. (f) Quantification of changes in ratio measured in d. All constructs were directly tagged with mCherry. *p<0.05, **p<0.01 ***p<0.001 Kruskal-Wallis Anova with Dunn’s multiple comparisons test. Data was obtained from three technical from three biological replicates (129< n <708) and is shown as mean±s.e.m.

**Supplementary Figure 6.**
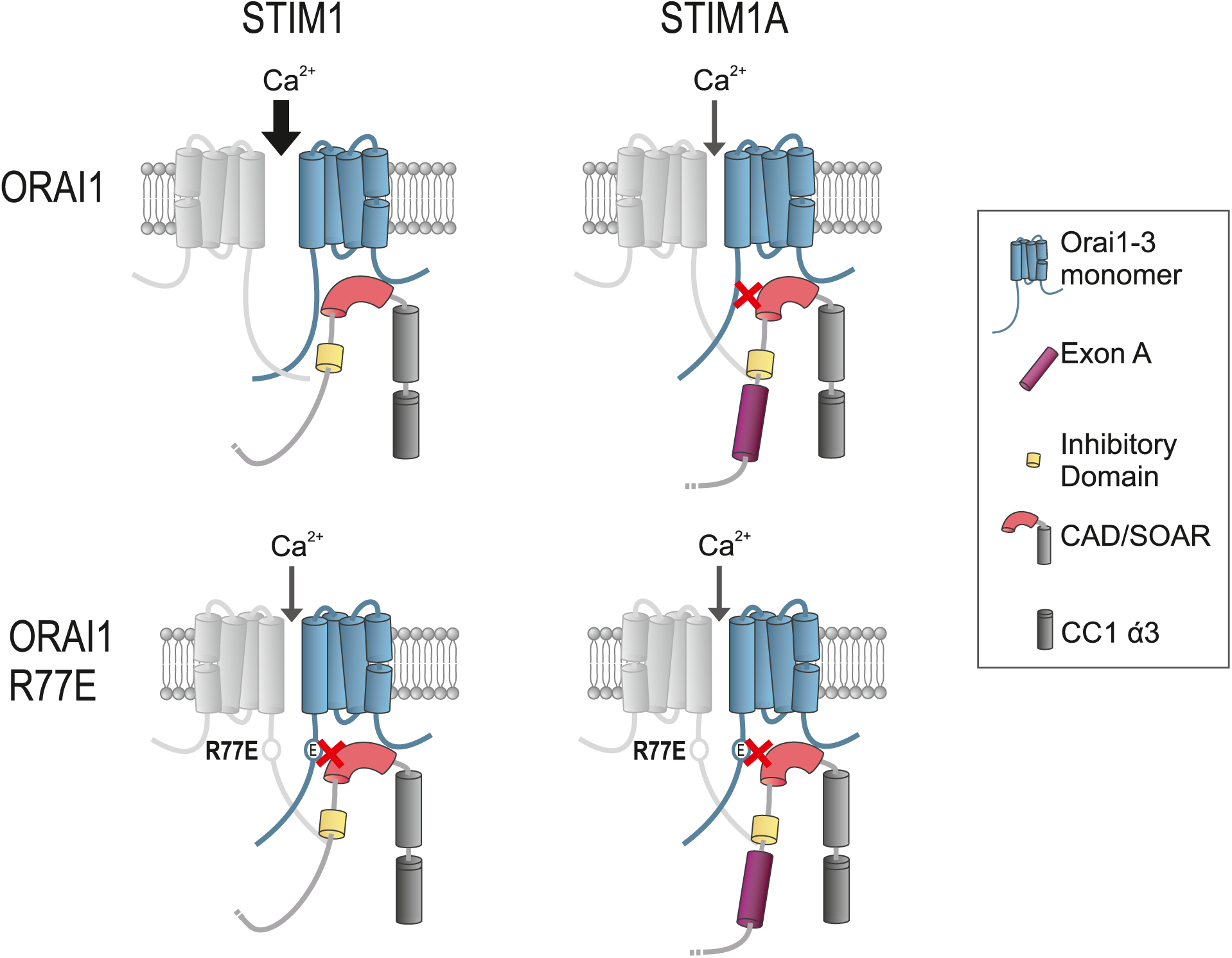
Model of the potential gating mechanism of STIMlA. Parts of the different cytosolic domains of STIMI, STIMIA and STIMIA D504A are shown as cylinders (CAD red/grey, ID yellow, exon A purple). Orai monomers are illustrated in light blue, as well as light grey. Orai point mutation R77E is labeled as a circle. STIMIA point mutation D503A is shown as a kink in exon A. The amount of Ca^2+^ entering the cell is illustrated as an arrow.

**Supplementary Table 1.** DNA-constructs.

| Construct Name | GOI | Tag | Vector |
| --- | --- | --- | --- |
| Lonza GFP transfection vector | GFP |  | pmax |
| mStim1_pmax_IRES_mcherry | Stim1 1 | IRES-mcherry | pmax |
| mStim1A pmax_IRES_mcherry | Stim1A | IRES-mcherry | pmax |
| mStim1A_pCAGGS-IRES_GFP | Stim1A | IRES-GFP | pCAGGS |
| mStim1_pCAGGS-IRES_GFP | Stim1 | IRES-GFP | pCAGGS |
| mOrai1_pmax_IRES_GFP | Orai1 | IRES-GFP | pmax |
| mOrai2_pmax_IRES_GFP | Orai2 | IRES-GFP | pmax |
| mOrai3_pmax_IRES_GFP | Orai3 | IRES-GFP | pmax |
| STIM1_C-term_mCherry | STIM1 | N-terminal HA, C-term_mCherry | pmax |
| STIM1A_C-term_mCherry | STIM1 | N-terminal HA, C-term_mCherry | pmax |
| STIM1A D503A_C-term_mCherry | STIM1 | N-terminal HA, C-term_mCherry | pmax |
| STIM1A S502A_C-term_mCherry | STIM1A | N-terminal HA, C-term_mCherry | pmax |
| STIM1A S502D_C-term_mCherry | STIM1A | N-terminal HA, C-term_mCherry | pmax |
| STIM1A RRFDS to AAAAA_C-term_mCherry | STIM1A | N-terminal HA, C-term_mCherry | pmax |
| STIM1A DDQS to AAAA_C-term_mCherry | STIM1A | N-terminal HA, C-term_mCherry | pmax |
| STIM1A FSD to AAA_C-term_mCherry | STIM1A | N-terminal HA, C-term_mCherry | pmax |
| STIM1A RR to QQ_C-term_mCherry | STIM1A | N-terminal HA, C-term_mCherry | pmax |
| pEX-YFP-hSTIM1 | STIM1 | N term YFP | pEX |
| HA_CRACM1_pCAGGSM2-IRESGFP | ORAI1 R77E | IRES GFP | pCAGGSM2 |
| HA_CRACM1_R77E/pCAGGSM2-IRESGFP | ORAI1 R77E | IRES GFP | pCAGGSM2 |

**Supplementary Table 2.**
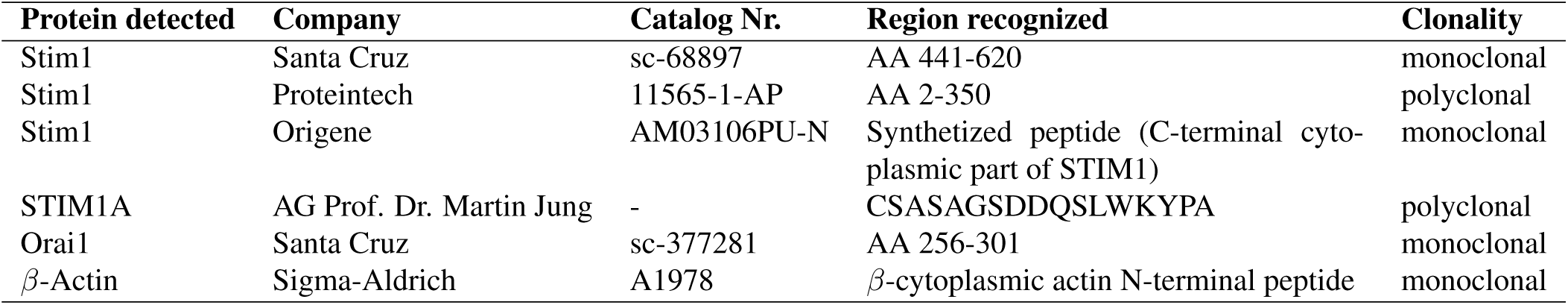
Antibodies.

**Supplementary Table 3.**
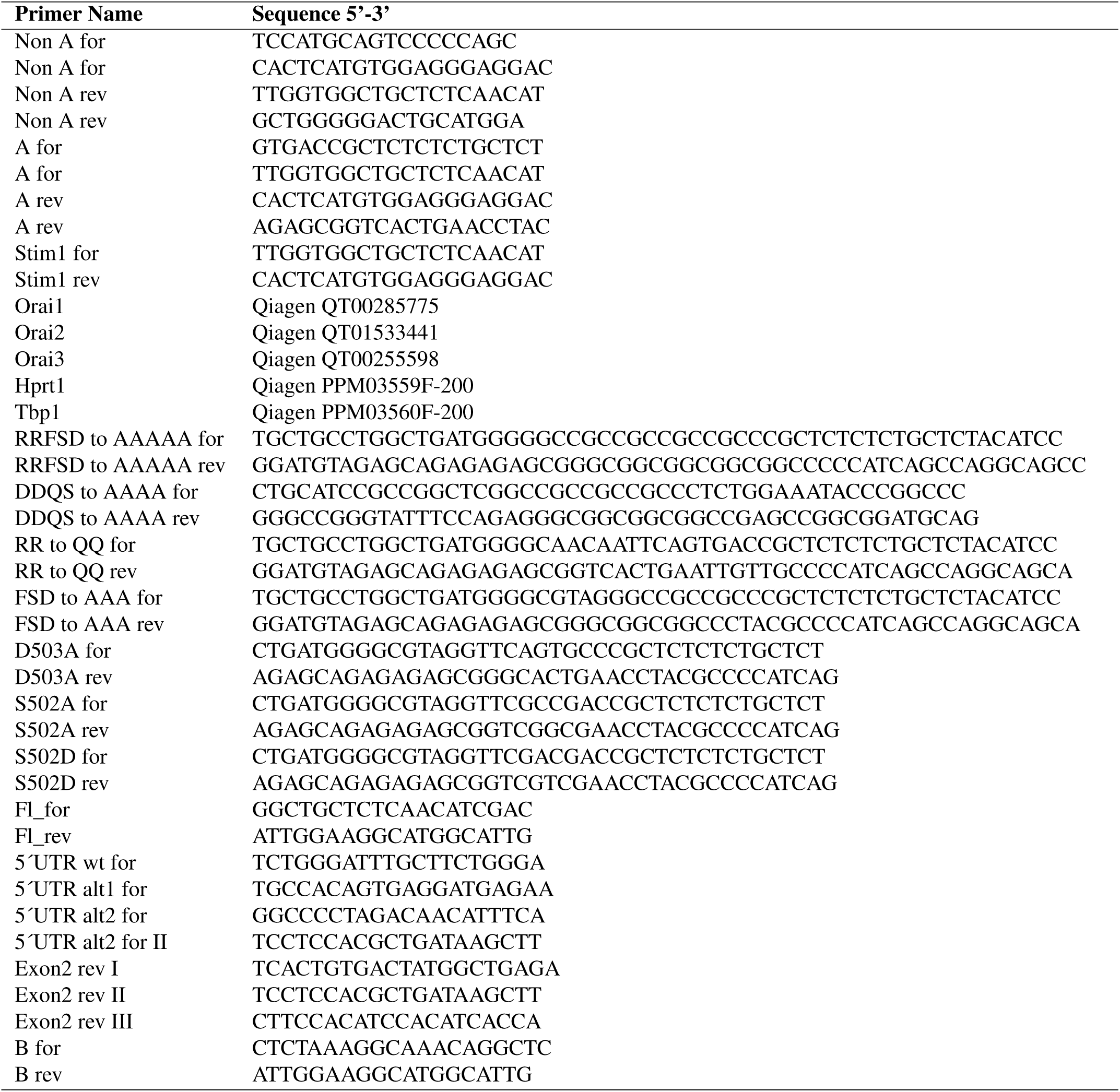
Primer.

